# PMPNN-DDG: an accurate machine learning-based ΔΔ*G* prediction pipeline trained on a novel interpretable feature set extracted from ProteinMPNN

**DOI:** 10.64898/2026.08.23.746499

**Authors:** Sajid Ahmed, Md Rafsan Jani

**Author notes:** At the time the experiments were conducted, Sajid Ahmed was a PhD student at Vanderbilt University and Md Rafsan Jani was a PhD student at Drexel University. These authors contributed equally to this work as co-first authors. Both authors are currently independent researchers.

## Abstract

An accurate and tractable approximation of the single-point mutation-induced change in protein thermodynamic stability, denoted by DDG, is critical for understanding the genotype– phenotype relationship. Several computational methods have been proposed for this problem; however, limited and error-prone training data and the difficult-to-predict magnitude of structural perturbations make this a challenging task. Consequently, the computational predictors proposed throughout the past decade incrementally improved prediction performance by proposing novel features, combining existing features, task-adapted neural network architectures, loss functions, data augmentation techniques, and pre-training procedures. In this work, we propose PMPNN-DDG, a Random Forest-based DDG prediction model, trained on a novel set of interpretable features extracted from the recently proposed message-passing neural network-based fixed backbone protein design model, ProteinMPNN. On the S_669_ independent test set, PMPNN-DDG achieves *r*_*F* +*R*_ = 0.64 and RMSE = 1.45, outperforming all compared baseline methods across the reported evaluation measures. On the S_sym_ independent test set, it achieves *r*_*F* +*R*_ = 0.81, *r*_*F−R*_ = −0.99, and RMSE = 1.10, showing competitive performance relative to the compared baselines. PMPNN-DDG is publicly available at https://github.com/dRanger666/PMPNN-DDG.

## 1 Introduction

Protein structural stability is a critical factor for successful protein folding and thus appropriate function [1, 2]. Consequently, single-point mutations that alter a protein’s amino acid sequence can disrupt folding and function through perturbations of thermodynamic stability [3]. Experimental quantification of this perturbation by measuring the folding-unfolding Gibbs free energy change poses substantial time and resource constraints, especially for high-throughput characterization. To address these issues, accurate and computationally efficient prediction pipelines are therefore needed.

Throughout the past decade, several computational pipelines have been proposed for predicting the impact of single-point mutations on protein stability [4]. Many of these methods use fast, lightweight machine-learning (ML) models that learn a mapping from mutations to corresponding DDG (i.e., ΔΔ*G*) values. A comparison of these pipelines [4] identified convolutional-neural-network (CNN)-based ACDC-NN [5], random-forest (RF)-based PremPS [6], 3D-CNN and atom-level-voxelization-based ThermoNet [7], untrained linear-feature-combination-based DDGun3D [8], and Rosetta-based energy-difference estimation [9] as top-performing methods in that benchmark.

The training datasets generally contain more instances of destabilizing mutations (DDG *>* 0) compared to stabilizing ones (DDG *<* 0). This results in significant prediction bias towards destabilizing variants. For mitigating this issue, ACDC-NN, PremPS, ThermoNet, and a few other recently proposed predictors have included reverse instances for each of the mutations in the training set, thus increasing the size of the training sets and removing skewness of the experimental label distribution. This augmentation has been shown to be effective in enforcing the highly desirable antisymmetric property [10] of these predictors while improving overall prediction performance. However, we argue that since the structural features of PremPS are extracted based on residue-level spatial information [6], values across those dimensions would be very similar for corresponding forward and reverse mutant pairs. This will result in potentially low information content of those structural features when reverse-mutant-based augmentation is applied. This could be one of the issues behind relatively low importance ranking of these features compared to the sequence-based evolutionary features in PremPS [6].

On the other hand, ACDC-NN [5] marginally outperforms its solely sequence-based counterpart ACDC-NN-Seq [11]. ACDC-NN incorporates structural information through position-specific scoring-matrix (PSSM) row encoding based on the local spatial neighborhood of the mutated residue. This input representation may be highly similar between wild-type structures and corresponding mutant models, which are passed through a shared Siamese neural network. Although ACDC-NN is trained to learn the antisymmetric relationship between members of these pairs through a custom loss function, a more informative input embedding has the potential to improve overall prediction performance. Additionally, due to limited experimentally labeled instances and inherent labeling noise, ACDC-NN highlights the importance of transfer learning while training ML-based DDG predictors, and attempts to address this issue by pre-training on predictions from the untrained predictor DDGun3D [8].

In a similar line of work, Chu et al. utilized embeddings extracted from the BERT-style masked protein language model ESM-1b [13, 12] to train graph-neural network (GNN) models on deep mutational scanning (DMS) datasets [14]. A simple fully connected neural network trained solely on embeddings corresponding to the mutated position was shown to significantly outperform the structural-geometry aware GNN models. Based on this observation, we hypothesize that more innovative methods with underlying biophysical interpretation are required for extracting informative representations from NN models trained on large sequence and structure databases.

To address these limitations, we introduce PMPNN-DDG, which uses the message-passing encoder-decoder autoregressive protein design model, ProteinMPNN (PMPNN) [15] to extract a minimal set of interpretable features with significantly non-overlapping information content. These features, inherently capable of capturing the differences between members of forward-reverse pairs, can ideally quantify the structural frustration induced by single point mutations. Subsequently, these are combined with PSSM-derived evolutionary conservation features, before training a RF regressor model. On the reported benchmarks, the resulting model outperforms all compared methods on S_669_ and shows competitive performance on S_sym_.

## 2 Benchmark Datasets

In this study, we use one dataset for training and validation of the RF model and evaluate PMPNN-DDG on three independent test sets:

1. S_2648_, comprising 2,648 manually verified variants with experimentally measured DDG values compiled in [16] from the database presented in [17], is used for training and validation. The 2,648 mutations in this dataset span across 131 distinct proteins. We excluded one protein and its corresponding mutations as the respective PDB record is obsolete.
2. S_669_, which was presented in [4] for unbiased ranking of the existing DDG prediction methods. It contains 669 variants belonging to 94 proteins having no more than 25% sequence similarity with any protein in S_2648_. It was compiled after filtering from the recently presented database ThermoMutDB [18].
3. S_sym_, containing 684 variants with experimental structures, where half of them are forward mutations, and the rest of them are corresponding reverse mutations, was presented in [19].
4. S_921_, presented in [6], containing 921 mutations from 54 proteins, was used as the third independent test set for additional comparison.

The three independent test sets were kept separate from model training and hyperparameter tuning to support valid comparison with the benchmarked methods.

## 3 Description of ProteinMPNN-extracted Features

A novel contribution of this work is the ProteinMPNN-based feature-extraction method, which encodes each single-point mutation using a nine-dimensional real-valued vector (i.e., ℝ^9^). These nine features are denoted by A, B, C, D, E-1, E-2, E-3, E-4, and E-5 throughout the rest of this manuscript. The last five features can be grouped together as these are generated by the same function; thus, E will be used to denote these five numbers as a set. Before each of the five feature generation functions are detailed in individual subsections, our general procedure to identify most attended neighbors, mask the identified neighbors one by one, extract corresponding embedding vectors, corresponding message vectors, and corresponding probability distributions from ProteinMPNN for quantifying the shift in structural stability induced by wildtype-to-mutant residue-identity change, is explained in the following subsection.

### 3.1 Neighbor Ranking and Neighbor-Vector Extraction Procedure

We hypothesize that the changes in center-to-neighbor message vectors, and the changes in the neighbor embedding vectors due to the change in embedding of the center-node, reflect the structural stability shift due to a particular single-point mutation. This is since this specific mutation information provided to the ProteinMPNN decoder has caused the message and embedding vectors to change in the first place with an attempt to alleviate potential structural frustration. Consequently, we establish a protocol to systematically extract these vectors for both the wild-type and the mutant sequences, which is the basis of our features B, C, D, and E. This protocol is illustrated in Figure 2.

**Figure 1:**
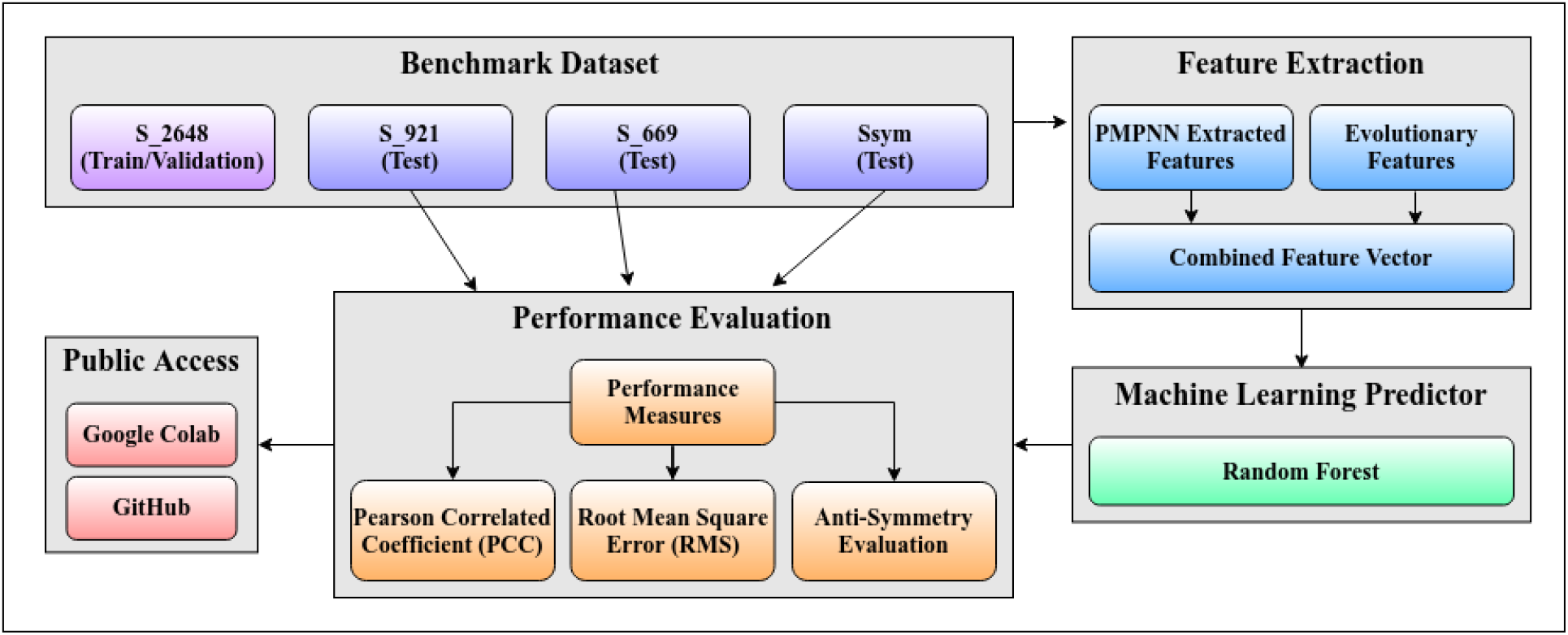
System diagram of PMPNN-DDG. We select one training/validation set (S) and three independent test sets from the literature, extract and analyze the proposed features from ProteinMPNN [15], combine them with evolutionary-conservation features, train a random-forest regression model on the combined feature vector, evaluate PMPNN-DDG using the reported performance measures, and compare it with methods included in the 2022 benchmark [4].

**Figure 2:**
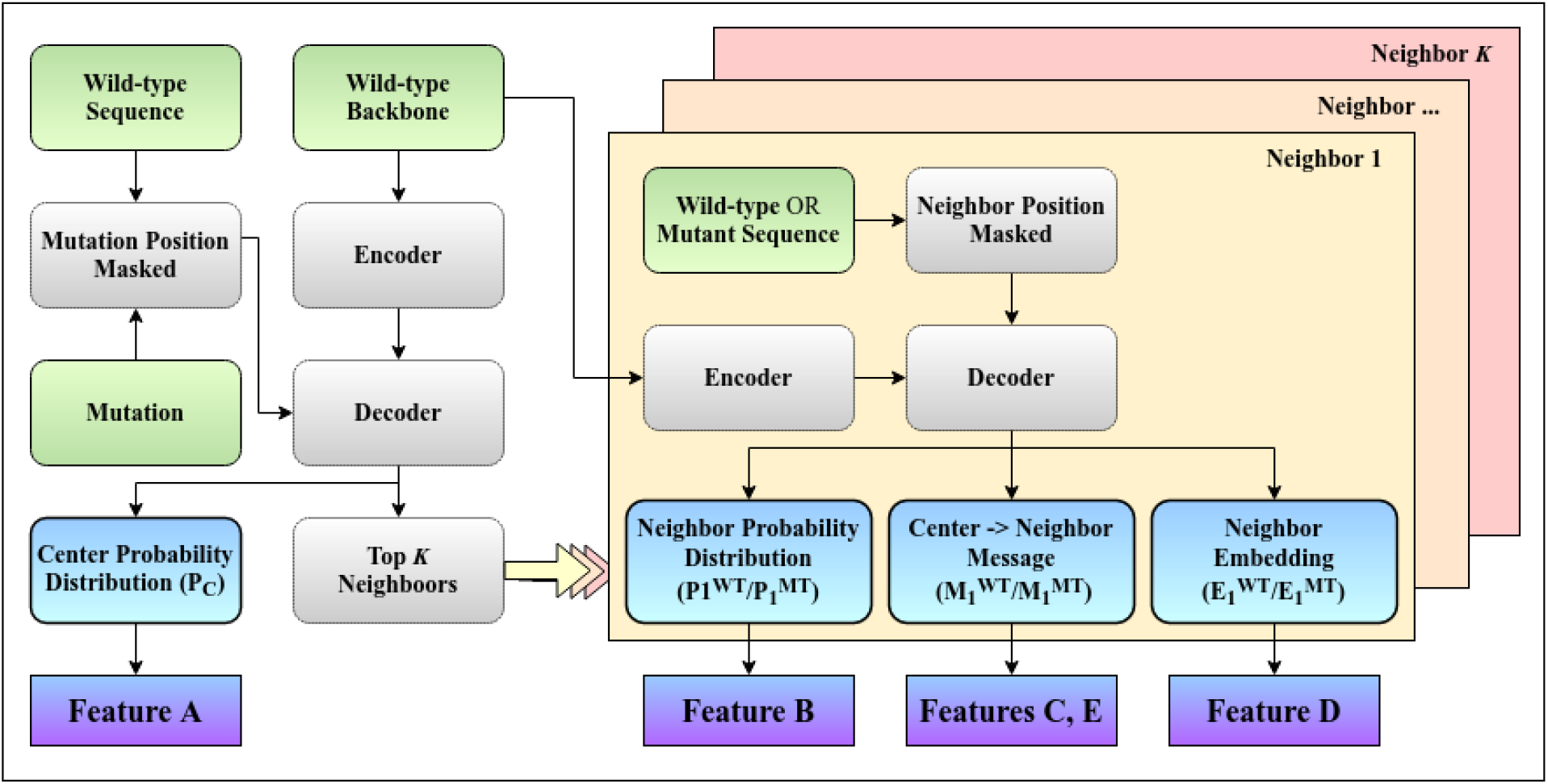
Overview of the proposed ProteinMPNN-based feature-extraction procedure. Input tensors are shown in green; ProteinMPNN encoder, decoder, and embedding sub-networks are shown in grey; output vectors extracted from ProteinMPNN are shown in blue; and the derived features are shown in violet.

As depicted in Figure 2, the first step is to pass the wild-type protein through ProteinMPNN, where the wild-type backbone is processed through the encoder layers before wild-type sequence embeddings are integrated at the decoder level. At this step, the mutation position is masked in the sequence input, thus ProteinMPNN attempts to reconstruct that position based on full backbone geometry and residue identity of all other positions, which are unmasked. The mutation position is referred to as the “center” or “center-node” throughout the rest of this section, since ProteinMPNN treats the input backbone as a graph, where every residue is connected to 48 nearest residues in three-dimensional space residues through directed edges, a hyperparameter used by ProteinMPNN. These 48 residues are referred to as “neighbors” or “neighbor-nodes”. Next, for each of these 48 neighbors, we extract a 128-dimensional vector, as defined by the ProteinMPNN architecture, that the center-node receives from the respective neighbor at the last decoder layer to update its own embedding (the “neighbor*→*center message vector”). Since each of these vectors can be regarded as a specific neighbor’s contribution to the center node’s identity, we utilize the *L*2-norms of these vectors to rank the neighbors based on their interaction strength with the mutated position in the wild-type structure, where a higher *L*2-norm would indicate stronger interaction). Subsequently, we select *K* neighbors with the highest norms, where *K* was set to 15 based on empirical observations. These neighbors, referred to as “Top *K* Neighbors” in Figure 2, can also be regarded as “the most attended neighbors” by the center node, since some hypothetical attention module’s coefficients would be used as scaling factors for the message vectors before summation [20]. Thus, we are approximating those coefficients by taking norms of those vectors as ProteinMPNN does not use any explicit attention module. The positions of these top-*K* neighbors are memorized at this point and utilized in the next step of our pipeline. Additionally, feature A is generated from the probability distribution corresponding to the masked center node extracted during this step of our pipeline (this point is discussed further in one of the following subsections, and the corresponding output vector is denoted by *P*_*C*_ in Figure 2).

In the next step, each of the top-*K* neighbor positions are masked one by one individually, while the mutation position is unmasked. The mutation position is set to both the wildtype residue and the mutant residue sequentially, which results in two sets of vectors (embedding vector, center*→*neighbor message vector, and probability distribution vector) for each neighbor position. One set corresponds to the wildtype center, and another set corresponds to the mutant center. The embedding vector, center*→*neighbor message vector, and probability distribution for neighbor *j* corresponding to wildtype center are denoted by 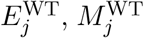 and 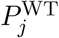, respectively, whereas the vectors in the same order for neighbor *j* corresponding to the mutant center are denoted by 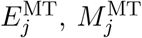, and 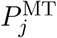, respectively. In the described setting, the differences between the equivalent vectors for the same neighbor position (i.e., 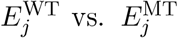, 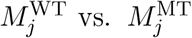 and 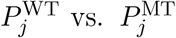) are conditioned on the change in center embedding due to WT *→* MT mutation. Therefore, we utilize these differences for characterizing mutation impact on the most attended neighbors. Afterwards, these vectors are encoded through the features B, C, D, and E.

### 3.2 Neighbor-Vector Encoding Methods

#### 3.2.1 Neighbor Entropy-Change Summation (B)

Feature B, depicting the change in information entropy [21] of the most attended neighbor position (i.e., top-ranked neighbor positions) probability distributions, can be expressed through Equation (1). Here, 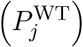 and 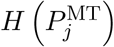 denote the information entropy values for the *j*^th^ neighbor corresponding to the wildtype center and the mutant center, respectively.

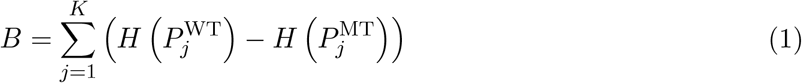

#### 3.2.2 Center-to-Neighbor Message-Norm-Ratio Summation (C)

Feature C summarizes the relative change in the magnitudes of center*→*neighbor message vectors induced by the WT *→* MT substitution and is defined by Equation (2). Here, 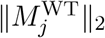 denotes the *L*_2_ norm of the message vector passed from the wild-type center to the *j*^th^ neighbor, whereas 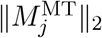 denotes the corresponding norm for the mutant center.

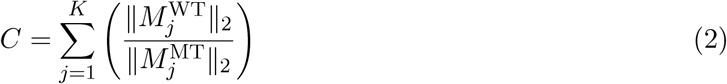

For feature C, we initially considered two schemes for encoding the change in each center *→*neighbor message vector induced by a WT *→* MT substitution: (1) the ratio of the two vector norms (NR) and (2) the norm of the corresponding difference vector (CN). As shown in the green subplot of Figure 3, NR has a higher correlation with the training labels than CN for feature C. We therefore use NR in Equation (2).

**Figure 3:**
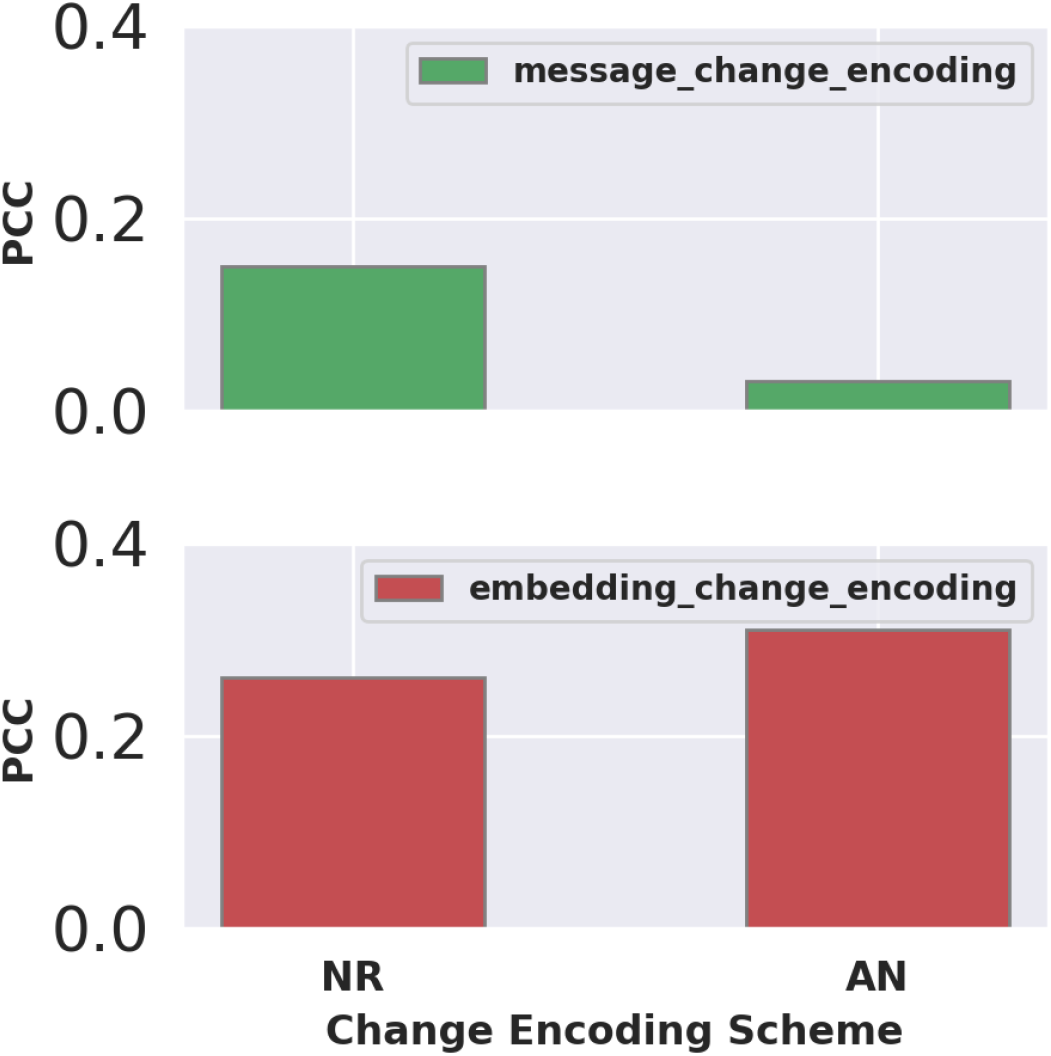
Comparison of norm-ratio (NR) and change-norm (CN) encoding schemes. PCC denotes the Pearson correlation coefficient between the proposed features C (green) and D (red) and the experimental ΔΔ*G* labels in the S_2648_ training set. For C, PCC is higher when NR rather than CN is used to compress each center*→*neighbor message change into a scalar. For D, PCC is higher for CN than for NR. We therefore use NR for C and CN for D.

#### 3.2.3 Neighbor Embedding-Change-Norm Summation (D)

Feature D, quantifying the change in the neighbor embedding vectors induced by WT*→*MT mutation, can be expressed as Equation (3). Here, 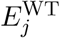 and 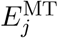 denote the *j*^th^ neighbor embedding corresponding to the wildtype and mutant centers, respectively. 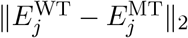 denotes *L*2-norm of the corresponding difference vector.

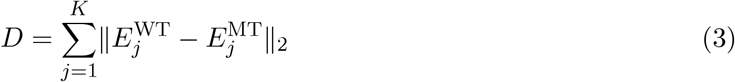

Encoding schemes NR and CN were tested for feature D as well. As shown in the red subplot of Figure 3, in case of D, CN is more correlated with the training labels compared to NR. Therefore, we use CN in Equation (3).

#### 3.2.4 Center-to-Neighbor Message-Change Kernel-PCA Projection Summation (E)

Feature E, encoding further discriminatory information regarding mutation structural stability impact from the changes in center*→*neighbor message vectors, can be expressed through Equation (4). Here, 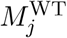 and 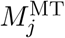 denote the *j*^th^ center*→*neighbor message corresponding to the wildtype and mutant centers, respectively. RBF_1*−*5_ denotes the transformation into the first five component scores returned by radial-basis-function (RBF) kernel PCA.

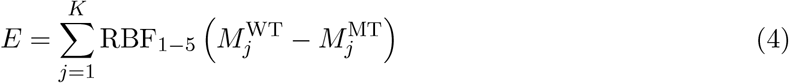

It is to be noted that, unlike B, C, and D, where multi-body energy changes are approximated using scalar values, E returns a length-five vector (the elements of this vector are denoted by E-1, E-2, E-3, E-4, and E-5 in a following section), since we have projected the center*→*neighbor message difference vectors onto the basis spanned by five leading principal components returned by a non-linear radial basis function (RBF) kernel-based PCA [22]. This kernel PCA was trained on randomly selected neighbor positions of mutations from the S_2648_ training set for computational tractability.

### 3.3 Mutation-Position Log-Probability Ratio (A)

The probability distribution corresponding to the mutation position (denoted as *P*_*C*_ in Figure 2), extracted during the first step of our pipeline, is utilized for generating feature A. Feature A, approximating mutation-position one-body energy change, can be expressed as Equation (5). Here, *P*_*C*_[WT] and *P*_*C*_[MT] denote the wild-type residue probability and the mutant residue probability at the mutation position, respectively.

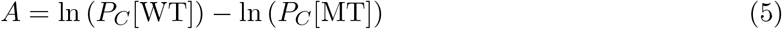

### 3.4 Feature-label and Pairwise Feature-correlation Analysis

While extracting the features A, B, C, D, and E from ProteinMPNN one by one, we attempt to get an initial estimation of how potent these individually are for the task of experimental DDG prediction, and how they might perform together when combined through a downstream ML model. For this purpose, we examine the Pearson Correlation Coefficient (PCC) of each of the features with experimental DDG labels along with the pairwise feature-feature PCC values on the S_2648_ training set, which are reported through the correlation heatmap in Figure 4.

**Figure 4:**
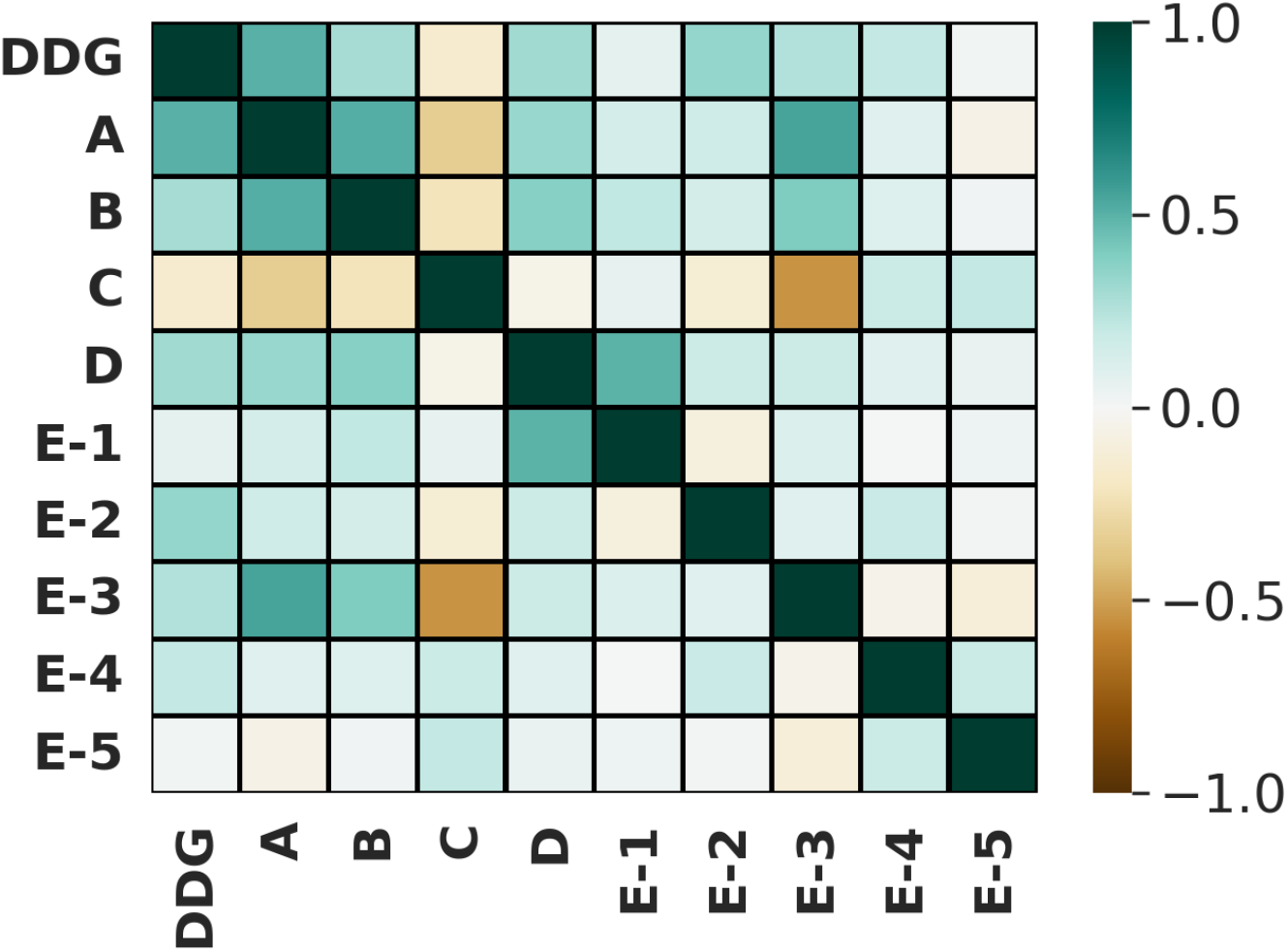
ProteinMPNN-extracted feature-feature correlation heatmap for S_2648_ training instances. DDG refers to experimental DDG labels.

As can be seen from Figure 4, among A, B, C, and D, all pairs seem sufficiently uncorrelated, for example, the maximum pairwise PCC of 0.52 is observed between A and B. On the other hand, feature C shows a PCC of -0.15 with the corresponding experimental DDG values, which is the minimum feature-label correlation in terms of PCC magnitude.

Consequently, some interesting patterns emerge after PCC values corresponding to E-1, E-2, E-3, E-4, and E-5 are included in the heatmap, which provide insight into the information captured by the PCA components, and their potential contribution to downstream ML model. E-1 shows a PCC of 0.49 with D, indicating that the maximum variation in center*→*neighbor message difference vectors could be largely explained by the changes in corresponding neighbor embedding vectors. This is somewhat intuitive since some amount of neighbor embedding change can be attributed to last decoder layer center*→*neighbor message change. On the other hand, E-2, which has the highest PCC magnitude with experimental labels among the five PCA-based features, is not significantly correlated with any other feature, featuring a maximum PCC of 0.17 with D), thus making it highly probable that incorporation of E-2 will boost downstream ML model’s performance. Similarly, E-4 has a PCC of 0.21 with the labels but has a maximum PCC of 0.20 with other features, which makes it another great candidate for incorporation into the combined feature set. While E-5 has low PCC with both the label column and the other features, E-3 has PCC values of 0.55, 0.40, and 0.55 with A, B, and C, respectively, and PCC of 0.26 with the label column. Features A, B, and C could probably account for this component if it were to be removed from the final feature set. Furthermore, E-1, E-2, E-3, E-4, and E-5 seem sufficiently decorrelated among themselves, for example, the maximum pairwise PCC of 0.20 is observed between E-2 and E-4.

Thus, These observations state that our proposed attributes A, B, C, and D contain significant discriminatory information with inherent biophysical interpretation, while group E increases information content of the combined set by supplementing the feature space with dimensions where abstract aspects of mutation structural impact are captured.

## 4 Description of Evolutionary Features (F, G, and H)

Most of the top-performing sequence-based or structure-based DDG prediction tools compared in [4], including PremPS [6], ACDC-NN [5], DDGun3D [8], and INPS-Seq [10], explicitly incorporate mutation-induced evolutionary-conservation information. These information, extracted through PSI-BLAST [23], HMMER3 [24], or PROVEAN [25], is utilized either as manually extracted feature encoding, or as part of sequence embedding provided to neural network-based representation learning modules. These features have usually been ranked as the top contributors [6], thus indicating that the downstream ML models or neural network architectures are at some level learning to extract structural stability related signals from mutation-position and/or mutation-neighborhood evolutionary patterns. Since our PMPNN-DDG features A, B, C, D, and E, lack any explicit multiple sequence alignment (MSA)-derived information, we explore the possibility of embellishing our final feature space by adding three evolutionary conservation descriptors.

The evolutionary features are extracted from position-specific scoring matrices (PSSMs). We construct PSSMs from protein-specific homologous sequences retrieved by running PSI-BLAST [23] against UniRef90 [26], using the default parameters described in [6]. For a particular mutation, the three evolutionary features are extracted from the row corresponding to the PDB sequence position at which the mutation has occurred and the two columns of that row containing the log-odds ratios corresponding to the occurrences of the wildtype residue and the alternate residue at the mutation site. Wildtype and mutant residue frequencies for the mutated position are referred to as *F*_WT_ and *F*_MT_, respectively. Our three evolutionary features, referred to as F, G, and H, denote (*F*_WT_ *− F*_MT_), *F*_WT_, and *F*_MT_, respectively.

Finally, we perform pairwise correlation analysis on the full feature set (A, B, C, D, E, F, G, and H). This analysis provides us with an estimation of how the evolutionary features (F, G, and H) might embellish the already informative feature space spanned by the ProteinMPNN-derived features (A, B, C, D, and E). The resulting correlation heatmap is shown in Figure 5.

**Figure 5:**
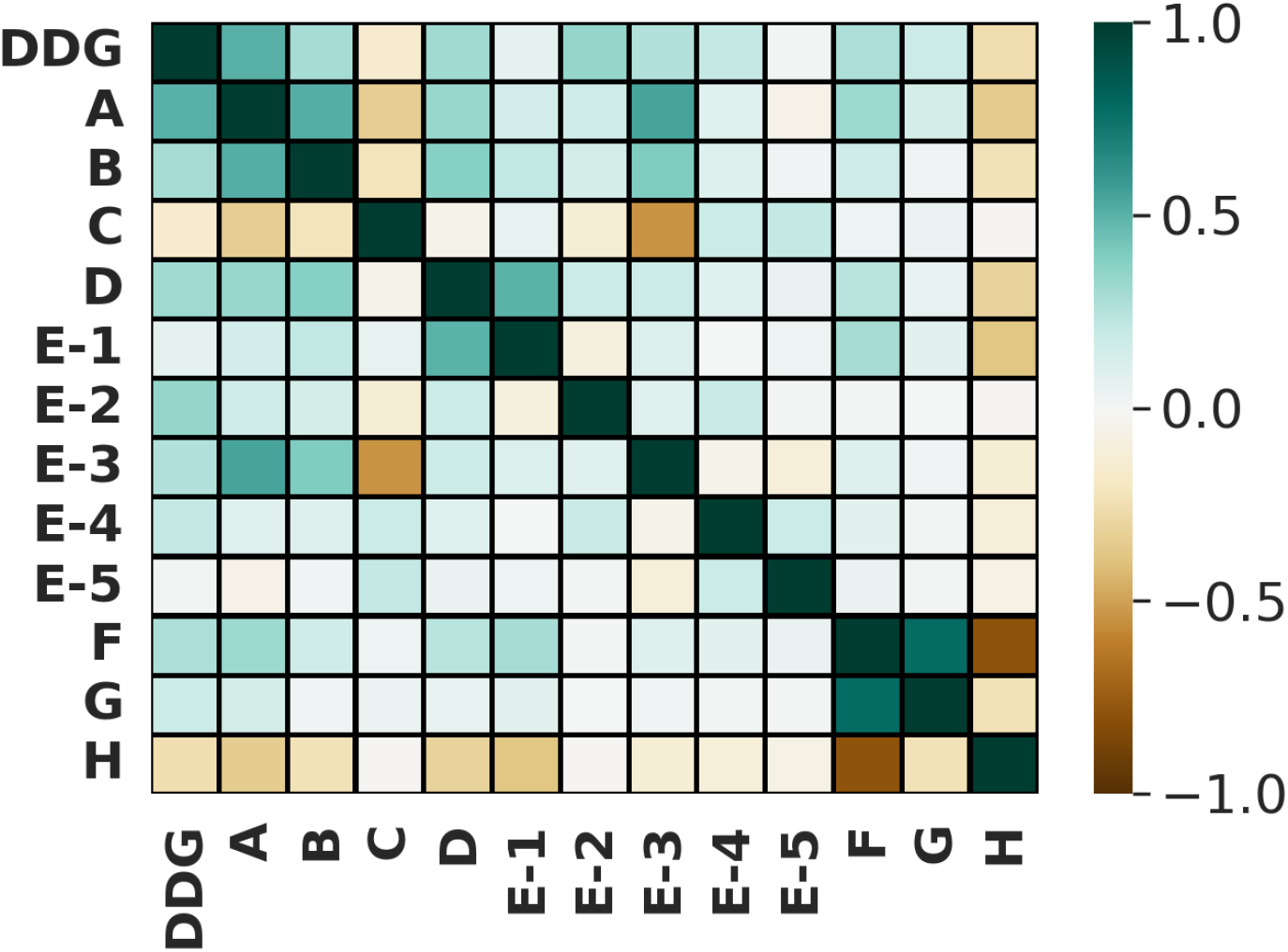
Final feature set correlation heatmap for S_2648_ training instances. DDG denotes the experimental labels. A, B, C, D, and E are the proposed ProteinMPNN-derived features, whereas F, G, and H are evolutionary features. The heatmap reports feature-to-label and pairwise feature-to-feature Pearson correlation coefficients.

As shown in Figure 5, features F, G, and H show maximum PCC magnitudes of 0.32, 0.15, and 0.39 with A and E-1, respectively. This indicates that most of the dimensions in our final feature space (the space spanned by ProteinMPNN-derived features + evolutionary features) contain significantly non-overlapping information encoding different aspects of the mutation under consideration. Having said that, it is to be re-iterated that E-3 might be removed without decreasing performance of the downstream ML model since the variation along E-3 is explained by A, B, and C i.e., some linear combination of these three features might recapitulate E-3 with substantially high accuracy. E-1 might also be removed due to its PCC magnitudes with both D and F. However, for this work, we choose to retain all five of the kernel-PCA projection features, and plan to address this issue with an attempt to improve downstream ML model performance further as part of future work.

## 5 Regression Model

The primary focus of this work is to setup a workflow that can extract a small non-redundant set of informative and interpretable features from ProteinMPNN, combine those with discriminative and “sufficiently” decorrelated evolutionary features, and establish the efficacy of the final feature set in predicting experimental DDG values against methods included in the benchmark of Pancotti et al. [4]. We use a Random Forest (RF) regressor [27], implemented in the scikit-learn [28] for mapping the feature vectors to experimental DDG values.

We performed a coarse-grained manual tuning for `n_estimators` and `max_samples` hyperparameters on held-out validation splits composed of all mutations from 25 randomly selected proteins from S_2648_, while training on mutations from the rest of the S_2648_ proteins. We selected `n_estimators` to be 500 and `max_samples` was set to 0.5 based on these validation results, while default scikit-learn values for all other hyperparameters were retained. We acknowledge that our predictor’s performance could improve further if a grid search-based extensive hyperparameter tuning was performed on more methodically held out validation partitions [29]. However, those empirical analyses are out of scope of this manuscript.

## 6 Performance Measures

We use several widely adopted measures to evaluate PMPNN-DDG on the independent test sets and to compare it with the methods evaluated in [4]. These are:

- Pearson correlation coefficient (PCC) between model predictions and experimentally determined DDG values.
- Root-mean-square error (RMSE) between model predictions and experimental DDG values.
- PCC between predictions for forward mutations and their corresponding reverse mutations, used as a measure of prediction antisymmetry.

## 7 Results and Discussion

### 7.1 Feature Contributions

As shown in Figure 6 for the independent test sets S_669_ and S_sym_, adding features A, B, C, D, E, F, G, and H one by one during training the RF model on S_2648_ training instances does not result in performance decrease between any two consecutive feature additions. This indicates that our final feature set fails to result in excessive dimensionality-induced overfitting. Moreover, up to the first evolutionary feature F, each dimension addition results in performance improvement for both independent sets, thus corroborating our decorrelation analysis-derived hypothesis regarding non-overlapping information content of the dimensions in the final space. However, addition of the last two evolutionary features G and H does not seem to result in further improvement (more pronounced for S_669_), which is again in agreement with pairwise correlation values (G and H have PCC values of 0.78 and -0.79 with F, respectively). Nevertheless, we retain features G and H as they fail to interfere with the primary focus of this work. It is to be noted that, for each of the results reported in Figure 6, the RF model was trained ten times on S_2648_, and PCC values derived from test set predictions of those ten models were averaged.

**Figure 6:**
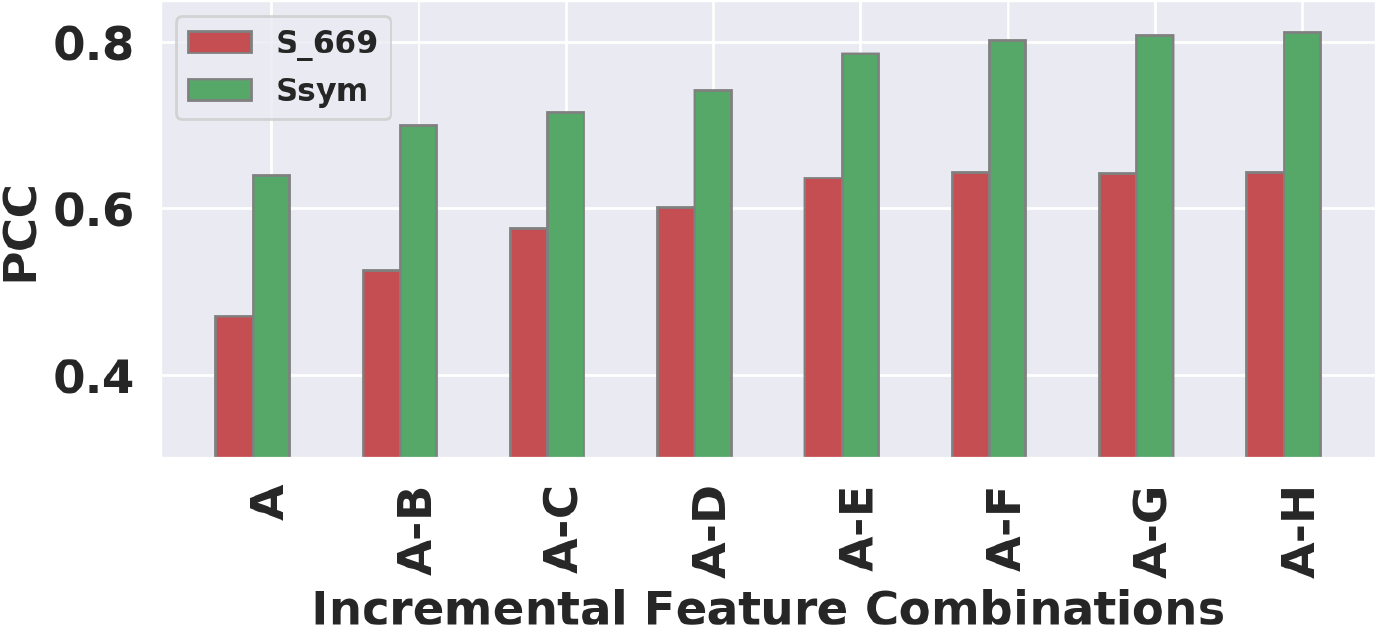
Incremental feature-combination analysis. Features are added cumulatively from A through H. The mean PCC on the S_669_ independent test set increases through the addition of F, whereas the mean PCC on the S_sym_ independent test set increases through the addition of G. In the figure labels, A–B, A–C, and so forth denote cumulative feature sets from A through the indicated feature.

### 7.2 Performance on Forward and Reverse Mutations and Antisymmetry of Predictions

Next, we investigate the performance of the RF model trained on our full feature set (A + B + C + D + E + F + G + H) on forward and reverse mutations of the test sets separately. Reverse mutations for S_2648_ instances were added during training as done in recent DDG prediction methods [4] for incentivizing the RF model to perform well on both stabilizing and de-stabilizing mutations. A PCC of -1 between forward and reverse predictions on the test sets would indicate that our predictor is perfectly antisymmetric. This is a highly desirable property for DDG predictors, since free-energy change introduced by a forward mutation plus the change induced by its corresponding reverse mutation should be equal to zero [10]. Additionally, since the majority of the forward mutations are in the first quadrant and most of the reverse mutations are in the third quadrant, correlation between predictions and experimental DDGs calculated on forward+reverse mutations might result in higher correlation compared to correlations on only forward and only reverse mutations [6]. Therefore, it is important to check only forward, only reverse, forward+reverse, and forward-vs.-reverse correlations for comprehensive evaluation of a DDG predictor. Only forward correlation, only reverse correlation, forward+reverse correlation, forward-vs.-reverse correlation, only forward root mean square error (RMS), only reverse RMS, and forward+reverse RMS, denoted by *r*_*F*_, *r*_*R*_, *r*_*F* +*R*_, *r*_*F −R*_, rms_*F*_, rms_*R*_, and rms_*F* +*R*_, respectively, are reported in Table 1.

**Table 1:** Performance of PMPNN-DDG on S_669_, S_sym_, and S_921_ independent test sets.

| Dataset | $r_F$ | $r_R$ | $r_{F+R}$ | $r_{F-R}$ | $\text{rms}_F$ | $\text{rms}_R$ | $\text{rms}_{F+R}$ |
| --- | --- | --- | --- | --- | --- | --- | --- |
| S<sub>669</sub> | 0.48 | 0.48 | 0.64 | -0.99 | 1.45 | 1.45 | 1.45 |
| S<sub>sym</sub> | 0.72 | 0.72 | 0.81 | -0.99 | 1.10 | 1.10 | 1.10 |
| S<sub>921</sub> | 0.77 | 0.77 | 0.79 | -1.00 | 1.49 | 1.49 | 1.49 |

As shown in Table 1 and observed for every antisymmetric DDG predictor in [4], *r*_*F* +*R*_ is significantly higher than both *r*_*F*_ and *r*_*R*_ on all three independent test sets. Moreover, PCC and RMSE on forward and reverse mutations are equal, which indicates that PMPNN-DDG performs equally well for stabilizing and destabilizing mutations, and *r*_*F* − *R*_ of -0.99 points towards the impressive antisymmetric property of model predictions.

### 7.3 Comparison with Methods Evaluated in the 2022 Benchmark

We next compare PMPNN-DDG with the ΔΔ*G* prediction methods evaluated by Pancotti et al. [4]. Results on the S_669_ dataset are presented in Table 2.

**Table 2:** Performance comparison of PMPNN-DDG with methods evaluated in the 2022 benchmark on the S_669_ independent test set.

| Method | $r_F$ | $r_R$ | $r_{F+R}$ | $r_{F-R}$ | $RMSE_F$ | $RMSE_R$ | $RMSE_{F+R}$ |
| --- | --- | --- | --- | --- | --- | --- | --- |
| **PMPNN-DDG** | **0.48** | **0.48** | **0.64** | **-0.99** | **1.45** | **1.45** | **1.45** |
| ACDC-NN | 0.46 | 0.45 | 0.61 | -0.98 | 1.49 | 1.50 | 1.50 |
| ThermoNet | 0.39 | 0.38 | 0.51 | -0.85 | 1.62 | 1.66 | 1.64 |
| PremPS | 0.41 | 0.42 | 0.62 | -0.85 | 1.50 | 1.49 | 1.49 |
| DDGun3D | 0.43 | 0.41 | 0.57 | -0.97 | 1.60 | 1.62 | 1.61 |
| Rosetta | 0.39 | 0.40 | 0.47 | -0.72 | 2.70 | 2.68 | 2.69 |

As shown in Table 2, PMPNN-DDG outperforms all compared baseline methods on S_669_ for every reported evaluation measure. Among the baseline methods, PremPS [6] has the highest *r*_*F* +*R*_, whereas ACDC-NN [5] and DDGun3D [8] have forward-vs.-reverse correlations closer to *−*1 than the other baselines. We next compare PMPNN-DDG with the same methods on S_sym_ in Table 3.

**Table 3:** Performance comparison of PMPNN-DDG with methods evaluated in the 2022 benchmark on the S_sym_ independent test set.

| Method | $r_F$ | $r_R$ | $r_{F+R}$ | $r_{F-R}$ | $RMSE_F$ | $RMSE_R$ | $RMSE_{F+R}$ |
| --- | --- | --- | --- | --- | --- | --- | --- |
| PMPNN-DDG | 0.72 | 0.72 | 0.81 | **-0.99** | 1.10 | **1.10** | **1.10** |
| ACDC-NN | 0.62 | 0.60 | 0.70 | -0.98 | 1.46 | 1.51 | 1.48 |
| ThermoNet | 0.45 | 0.39 | 0.50 | -0.89 | 1.66 | 1.73 | 1.70 |
| PremPS | **0.81** | **0.73** | **0.84** | -0.93 | **1.05** | 1.21 | 1.14 |
| DDGun3D | 0.58 | 0.56 | 0.66 | **-0.99** | 1.45 | 1.49 | 1.47 |
| Rosetta | — | — | — | — | — | — | — |

As shown in Table 3, PremPS has higher *r*_*F*_, *r*_*R*_, and *r*_*F* +*R*_ and lower RMSE_*F*_ than PMPNNDDG on S_sym_. In contrast, PMPNN-DDG has a forward-vs.-reverse correlation closer to *−*1 and lower RMSE_*R*_ and RMSE_*F* +*R*_. Thus, the comparison on S_sym_ is mixed across the reported metrics.

### 7.4 Scope of the Comparison

The experiments and baseline comparisons reported in this manuscript were completed by October 2022. Accordingly, the comparison set reflects methods included in the 2022 benchmark and does not include methods introduced subsequently.

## 8 Conclusion

In this work, we present PMPNN-DDG, a machine-learning based computational pipeline for estimating single-point mutation-induced shift in protein thermodynamic stability. The model comprises a random forest regressor trained on a feature set containing nine novel interpretable structural features extracted from protein design neural network ProteinMPNN, and three evolutionary features. On S_669_, PMPNN-DDG outperforms all compared baseline methods across the reported metrics. On S_sym_, performance is mixed relative to PremPS: PMPNN-DDG shows stronger antisymmetry and lower combined RMSE, whereas PremPS achieves higher forward, reverse, and combined correlations.

In fact, this work introduces an innovative paradigm for extracting informative interpretable features quantifying mutation-induced thermodynamic stability shift from a well-accepted protein design model [15]. The proposed feature extraction principles can be applicable to a variety of pretrained message-passing or attention-based neural networks, in general. Furthermore, the proposed features can be readily combined with non-overlapping information containing attributes that might be extracted from attention-based sequence and structural language models (i.e., evolutionary scale language models proposed in [12, 30]) through fundamentally similar mechanisms. We intend to explore this possibility as part of future work.

